# Spatial and depth structuring predominate over temporal variation in Mediterranean grassland soil viral communities

**DOI:** 10.64898/2025.12.30.696869

**Authors:** Jane D. Fudyma, Petar Penev, Katerina Estera-Molina, Jordan Hoff, Steven J. Blazewicz, Jennifer Pett-Ridge, Joanne B. Emerson

## Abstract

Viruses have the potential to influence microbial community structure and elemental cycling in soils, but it remains unclear how these communities are distributed across space and time, which can shape how they respond to environmental change and impact ecosystem processes. Mediterranean grasslands, with their pronounced seasonal moisture fluctuations, offer an ideal system to examine viral biogeography. While previous studies hint at spatial structuring and moisture controls on viral communities, temporal responses to seasonal moisture shifts have not been comprehensively investigated *in situ*. Here, we generated 59 viromes and leveraged 89 metagenomes from two Mediterranean grasslands to measure soil viral communities across horizontal space (sampling zone), depth, and key seasonal stages of the Mediterranean water year (e.g. plant productivity, dry down and wetup). Sampling zone was the dominant driver of viral community composition in viromes, with time secondarily explaining variation in viral communities. In contrast, viral richness and DNA yields varied primarily across time. Spatial structuring also emerged in viruses recovered from metagenomes, with depth having the strongest effect, followed by sampling zone. Environmental variables and predicted host distributions partially explained these patterns, but substantial variation remained unaccounted for, suggesting a role for dispersal limitation. Overall, soil viral communities were primarily structured by spatial factors, with temporal and environmental influences acting secondarily, highlighting the importance of fine-scale spatial dynamics in understanding viral ecology. Future studies should explicitly examine the role of dispersal limitation and fine-scale host-environment interactions to fully resolve drivers of soil viral biogeography.

## Introduction

With abundances often comparable to those of prokaryotes [1, 2], viruses are increasingly recognized as key players in soil ecosystems [3]. Virus-induced cell lysis can release host nutrients and restructure microbial communities, altering microbial composition and biogeochemical cycles [4–7]. In marine systems, these processes drive substantial ecosystem impacts, contributing to the daily turnover of up to 40% of microbial cells and releasing up to 10^4^ Mg C per second [8–11]. In contrast, the ecological drivers of soil viruses remain poorly understood [3, 5], such as how viral communities are distributed across terrestrial landscapes and what environmental factors shape their composition and abundance. This baseline understanding of soil viral biogeography is essential for recognizing how viruses respond to perturbations and how this in turn could impact ecosystem processes.

The relative importance of environmental and spatiotemporal drivers in shaping soil viral communities in the field remains unclear. Across pronounced environmental gradients, such as distinct habitats [12–18] or strongly differing edaphic properties (e.g., wide ranges of soil moisture or pH) [19–21], viral communities are structured predictably, consistent with environmental filtering. When environmental gradients are subtle or patchy, viral communities can be shaped more strongly by spatial heterogeneity [22–24] and can even differ over meter-scale distances [23, 25], suggesting that dispersal limitation could be an important force whose effects emerge most clearly in the absence of overriding environmental change. Vertical structuring has also been observed, with differences in viral community composition and virus-like particle (VLP) abundance across soil depths [15, 26], sometimes in alignment with edaphic changes [20]. Although longitudinal studies are limited, temporal shifts in soil viral community composition [13, 17, 22, 24, 27, 28] and VLP abundances [14, 29] have been observed. These temporal shifts tend to coincide with notable changes in environmental conditions, such as during snowmelt or post-fire [17, 24, 28], and/or occur in relatively spatially homogeneous, well-mixed systems, such as tilled agricultural soils [22]. Still, spatial variation sometimes explains more viral community variance than time, a pattern even observed over multiple years [27], consistent with stronger spatial than temporal structuring observed in soil prokaryotic communities [30–34]. Altogether, these findings reveal a complex, understudied viral biogeography, where the relative importance of environmental filtering and spatiotemporal gradients – especially across natural seasonal cycles in heterogeneous systems – remain unknown.

Mediterranean grasslands are an ideal system for disentangling the ecological forces that shape soil viral communities, as they exhibit strong, predictable environmental drivers (recurring seasonal shifts in precipitation) and have shown evidence of spatial structuring consistent with dispersal limitation [23]. These ecosystems undergo marked wet-dry cycles, with cool, wet winters followed by hot, dry summers [35]. As the dry season progresses and moisture rapidly declines, prokaryotic communities tend to change activity and lifestyle strategies, such as reducing growth or going dormant [35–39]. The first rains of the autumn season trigger microbial activity and increases in abundance [35, 36, 40], with microbial communities reorganizing rapidly in response [41–44]. Because viruses rely on microbial hosts, they too are expected to respond to these hydrological transitions, but existing results suggest viral community responses to soil moisture are inconsistent. Some microcosm studies, including one across multiple Mediterranean grassland soils, have observed blooms in viral diversity and abundance after rewetting [45, 46], while another microcosm study reported a decline in diversity with rewetting [7]. Together with suggestions of greater viral diversity in historically drier grasslands [47], some evidence of lower viral abundances in dry soils [3, 46], and the predominance of spatial distance over precipitation treatment in structuring viral communities [23], these mixed findings reveal a key knowledge gap: when strong temporal pulses in moisture occur within spatially heterogeneous field settings, do viral communities reflect temporal environmental change, underlying spatial structure, or some interaction of the two? Mediterranean grasslands offer a natural testbed to address this question and evaluate how viral communities are structured across space and time in response to environmental forcing.

Here we combined viromic and metagenomic approaches to examine how soil viral communities vary across spatial scales, soil depth, and moisture-driven seasonal stages of the Mediterranean water year (e.g., as captured in our study from March–November, spanning the dry-down and wet-up transitions). We focused on two grasslands that differ in historical precipitation and soil properties, allowing us to explore how local environmental conditions interact with spatial and temporal structuring. We hypothesized that spatial heterogeneity would be the primary factor shaping viral community composition, with temporal fluctuations modulating these patterns over the course of the water year.

## Materials & Methods

### Study sites & sampling

Soil samples were collected and processed as described previously [39]. Briefly, soil cores were taken at the Angelo Coast Range Reserve at South Meadow, CA, USA (Lat: 39.73925, Long: −123.6300) and the Hopland Research and Extension Center at Buck Field, CA, USA (Lat: 39.00178, Long: −123.0682, **Figure 1A**). Samples were collected at 10 time points from April-November 2022, spanning key phases of the Mediterranean water year: peak plant productivity (T1-T3), dry-down (T4, T5 (Hopland), T6 (Angelo), T7), and fall wet-up (T8 (Hopland), T9, T10 **Figure 1C**). At each timepoint, a core was taken from three sampling zones within each field, at a randomly determined point within one meter of the zone’s center (**Figure 1B**). Soil cores (7.6 cm diameter) were collected with a slide-hammer corer or a Geoprobe 54LT to 80 cm depths. Core inserts were sealed with sterile caps and parafilm, transported to the University of California, Berkeley (CA, USA), and subsampled for viromes and total metagenomes within the same day. Cores were first spit into five depth horizons: 0-10 cm, 10-20 cm, 20-30 cm, 30-50 cm, and 50-80 cm. Then for viromes, the 0-10 cm and 10-20 cm profiles were split vertically in half, and soils were sub-sampled from the interior of each half and homogenized together, creating two vertical virome replicates per core that spanned 0-20 cm depths. Virome soils were transported on ice to the University of California, Davis, CA, USA and kept at 4 °C until processing, within 24 hours. For metagenomes and soil chemistry, soils were processed by homogenizing and removing roots from the full core at each depth, with one replicate per depth per core. Soils for metagenomes were flash-frozen in liquid nitrogen and stored at −80 °C until processing.

**Figure 1:**
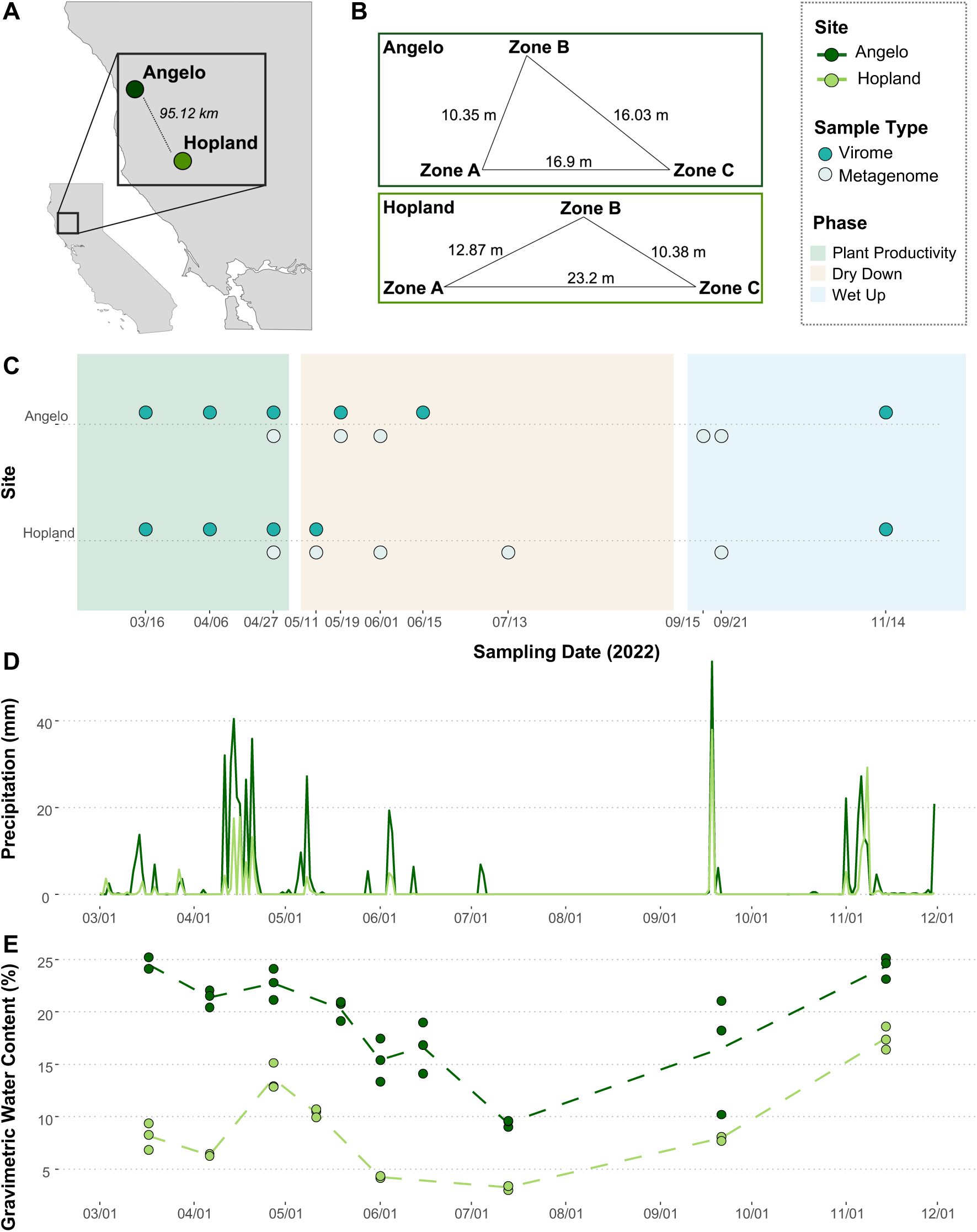
Experimental Design. **A)** Map showing the two Californian Mediterranean grasslands sampled in this study. **B)** Sampling design in each site, where at each timepoint, samples were collected within a ∼1-meter radius of the center of a designated sampling zone (zone A, B or C). Distances between each zone are denoted. **C)** Ten timepoints were sampled throughout key phases of the water year (early peak plant productivity, dry-down, and wet-up). Viromes and metagenomes were processed at different timepoints, as illustrated in the timeline plot. **D&E)** Daily average precipitation **(D)** and Gravimetric Water Content (**E** – GWC %) throughout the duration of the study. GWC was measured in triplicate, with one measurement for each sampling zone at a timepoint. Colors correspond to each site, with Angelo in dark green and Hopland in light green.

### Soil chemsitry

Gravimetric water content (GWC), pH, total carbon (C) and total nitrogen (N) were processed at University of California, Berkeley, USA, as described previously [39]. Briefly, GWC was measured via the oven-drying method, pH using CaCl_3_ (Oakton 110 series pH meter), and total N and total C with CHNOS Elemental Analyzer (varioISOTOPE cube, Elementar, Hanau, Germany) at the Center for Stable Isotope Biogeochemistry (University of California, Berkeley).

### Virome DNA extraction, library construction, and sequencing

Timepoints 1-4 and 10 (both sites) and 6 (Angelo) were processed for viromics. This sampling design was largely due to practical considerations (i.e., availability for processing viromes within 48 hours of collection; longer storage would have required freezing and different processing protocols [48]). VLPs were extracted as previously described [22, 48, 49]. For each sample, 9 mL of protein-supplemented phosphate buffered saline (PPBS) buffer was added to 10 g of soil, briefly vortexed, agitated for 10 min on an orbital shaker (300 rpm, 4 °C), centrifuged (10 min, 3,085 x g, 4 °C), and supernatant was poured into a new tube. This was repeated three times for a final volume of 27 mL of supernatant. The supernatant was centrifuged twice to remove residual soil particles (8 min, 10,000 x g, 4 °C) and subsequently filtered through a 0.22 µm filter. VLPs were pelleted by ultracentrifugation (2 h 25 min, 111,818 x g, 4 °C), in an Optima LE-80 K ultracentrifuge and 50.2 Ti rotor (Beckman Coulter Life Sciences). Pelleted VLPs were resuspended in 100 µL of UltraPure water and treated with Promega RQ1 DNase (1:10 ratio DNase, 30 min, 37 °C) to remove free DNA. Viral DNA was extracted using the DNeasy PowerSoil Pro kit (Qiagen) according to manufacturer’s instructions, with an additional heat treatment (10 min, 65 °C) before the bead-beating step. Libraries were constructed by the University of California, Davis DNA Technologies Core using the DNA KAPA HyperPrep library kit (Roche). Paired-end 150 bp sequences were generated using the NovaSeq 6000 (Illumina) to a depth of >10 Gbp per library.

### Metagenomic DNA extraction, library construction, and sequencing

Metagenomes were processed as described previously [39]. Samples were processed from five timepoints (3, 4, 5, 7(Angelo)/8(Hopland), 9) and three depths (0-10 cm, 20-30 cm, 50-80 cm). DNA was extracted from 10 g of soil using the QIAGEN DNeasy PowerMax Soil Kit, with a back-extraction modification step for deep soil profiles (>50 cm) to obtain enough DNA for sequencing. Libraries were prepared by Novogene, and paired-end 150 bp sequencing was performed on a NovaSeq 6000 platform (Illumina).

### Virome and metagenome quality control

Raw virome reads were quality filtered using Trimmomatic (v0.39, [50]), with a minimum quality-per-base score of 30 and a minimum read length of 50 bases. PhiX sequences were removed using BBduk from the BBMap package (v38.63, [51]). Metagenomic raw reads were processed as previously described [39]. Briefly, reads were assessed for quality with FastQC, llumina adapters were trimmed using BBduk, and phiX and Illumina artifacts were removed with BBduk. Metagenomic reads were then quality trimmed with sickle using default parameters.

### Virome assembly, viral species identification and host prediction

*De novo* assemblies were generated for each virome separately, using MEGAHIT (v1.0.6, [52]) in meta-large mode with a 10,000 bp minimum contig length requirement. Viral contigs were identified using VIBRANT (v1.2.1, [50]) with the virome flag, retaining only medium- and high-quality contigs. Contigs were dereplicated into a set of species-level viral operational taxonomic units (vOTUs) using dRep (v3.2.2,[53]), at 95% average nucleotide identity (ANI) with a 85% minimum coverage threshold, using the ANImf and single cluster algorithms. This dereplicated set of vOTUs was used for read mapping of both viromic and metagenomic reads, as well as host prediction. Host taxonomy was predicted using iPHoP (v1.2.1 [54]) with default parameters. If more than one host was predicted for a vOTU, we chose the prediction with the highest confidence score.

### Read mapping and abundance table generation

Viromic reads and, separately, metagenomic reads were mapped to the dereplicated vOTUs using Bowtie2 (v2.4.2,[55]). A trimmed-mean coverage table was generated for viromes using CoverM (0.6.1, [56]), with a 75% minimum breadth and a 90% minimum read identity. For metagenomes, since substantial non-viral sequences can make it difficult to recover vOTUs [22], we optimized read-mapping parameters to both enhance viral recovery and minimize non-viral false positives. We compared metagenomic read-mapping breadth of coverage thresholds from 0 to 75% (75% being standard for viromes) to paired viromes at timepoints 3 and 4, revealing that 5% coverage breadth for metagenomes generated the most similar vOTU relative abundances to paired viromes, and decreased false positives from no coverage threshold by 400-fold (**Figure S1**). Thus, we used 5% coverage breadth with 90% read identity to generate a metagenomic vOTU trimmed-mean abundance table, using CoverM. For viral richness analyses, due to differences in sequencing depth, each virome was rarefied by randomly subsampling the mapped reads by a factor calculated by dividing by the lowest sequencing depth from that of the sample, using SAMtools (v1.13, [57]), and a trimmed-mean table was generated using the same breadth and read percent identity as above.

### Statistical analyses

All data analyses were performed in RStudio (Version 2025.05.0+496, [58]) and tidyverse (v2.0.0, [59]). Bray-Curtis dissimilarities on Hellinger transformed relative abundances were calculated using vegan (v2.6.10, [60]). These were either visualized using Principal Coordinates Analyses (PCoA, ape, v5.8.1, [61]), or sample-to-sample pairwise comparisons as temporal distance-decay curves or boxplots (ggplot2, v3.5.2, [62]). PERMANOVA was performed on distance matrices (adonis2 – vegan, 999 permutations). UpSet plots were generated with ComplexUpset v1.3.3 [63], based on presence-absence data. Linear regressions and loess trends were fitted to temporal distance-decay curves, with goodness of fit determined by Pearson’s correlation coefficients. Either ANOVA or Dunn Tests (ggpubr, v0.6.0, [64]) were performed to determine significance of Bray-Curtis group variance by zone or sampling timepoint. For environmental and predicted host abundance comparisons by either sampling zone or timepoint, we fit linear models or generalized linear models (if linear model assumptions were not met) and determined significance by ANOVA, followed by pairwise post hoc tests (emmeans, v1.11.1, [65]). For distance-decay of environmental variables, we followed a similar approach as for viral community temporal distance-decay, but with environmental variables z-score transformed and Euclidean distances calculated for the distance matrix. Viral richness and viromic DNA yields were correlated using linear regressions. To determine richness and yield correlations with other variables (environmental, predicted hosts), we fitted loess trends to each variable throughout time and only retained variables that had moderate-to-strong Spearman’s rank correlations coefficients (R^2^ ≥ 0.5 or ≤ −0.5). We then took only strong time correlations and compared these to each other, again only reporting moderate-to-strong Spearman’s rank correlation coefficients. To correlate vOTUs with pH and total N (%) while controlling for depth, we fit generalized additive models (GAMs, mgcv, v1.9.3,[66]) for each vOTU using depth-corrected nitrogen and pH residuals. Significant relationships were identified based on FDR-corrected p-values. To assess the influence of soil environmental variables on richness of vOTUs recovered from metagenomic read-mapping, we used a linear mixed-effects model, with replicate included as a random effect (lme4, v1.1.37, [67]). All final plots were generated with ggplot2 and cowplot v1.3.3 [68].

## Results and Discussion

To investigate how seasonal variation in soil moisture affects soil viral diversity and compositional changes, we generated 59 viral size fraction (< 0.2 μm) metagenomes (viromes) and leveraged 89 total metagenomes from soils collected from two Mediterranean climate grasslands (Hopland and Angelo) throughout three key seasonal periods: peak plant productivity, dry down, and wet up (**Figure 1C**). Even though the sampling timeline was designed *a priori* based on historical precipitation data, we still largely captured the key changes in precipitation and soil moisture (**Figure 1D-E**), although late precipitation in Angelo prolonged the dry down period (**Figure 1D**). The two grasslands differed significantly in overall soil edaphic properties throughout the study (**Figure S2, Table S2**), consistent with their historically different temperature and precipitation regimes [69–72]. From the 59 viromes extracted from the top 0-20 cm from March-November, we recovered a total of 19,117 vOTUs (viral operational taxonomic units, species-level, [73]). From the 89 metagenomes taken from three depths (0-10 cm, 20-30 cm, and 50-80 cm) from May-October, we recovered 13,959 vOTUs from read-mapping to the virome database. We recovered 9,385 vOTUs from Angelo viromes, of which 7,718 were also recovered in Angelo metagenomic data via read mapping, and 10,764 vOTUs were recovered from Hopland viromes, of which 7,908 were detected in Hopland metagenomeic read mapping. The large number of vOTUs detected from metagenomic read mapping was attributed to (1) the use of a vOTU database for read-mapping generated from viromics, which enables greater vOTU assembly and detection by removing the bacterial and relic-DNA background that typically makes viral recovery difficult in total metagenomes [22, 74], and (2) the availability of 6 paired virome-metagenome samples, which enabled validation and relaxation of metagenomic read-mapping coverage thresholds (see Methods, and [19]). Any relaxation of recovery thresholds in future metagenomic analyses should be applied only with careful validation against viromic samples.

### Position within the field (sampling zone) best explained viral community composition in surface soils

In both field sites, viral community composition (measured via viromes) was primarily structured by sampling location within the field (collection from zone A, B, or C; PERMANOVA p < 0.001, R^2^ = 0.32 at Angelo, 0.24 at Hopland, **Figure 2A, Table S1**), with weaker contributions from sampling date (p < 0.001, R^2^ = 0.092 at Angelo, 0.12 at Hopland, **Table S1**). Although strong spatial structuring was evident at the field scale, samples collected in near proximity within the same core were most often the most similar to each other within the dataset (**Figure S3**). These patterns suggest that the strength of spatial structuring varies with scale (e.g., pronounced across the field but weaker between adjacent cores), likely reflecting the combined influences of local environmental variation and limits to viral dispersal.

**Figure 2:**
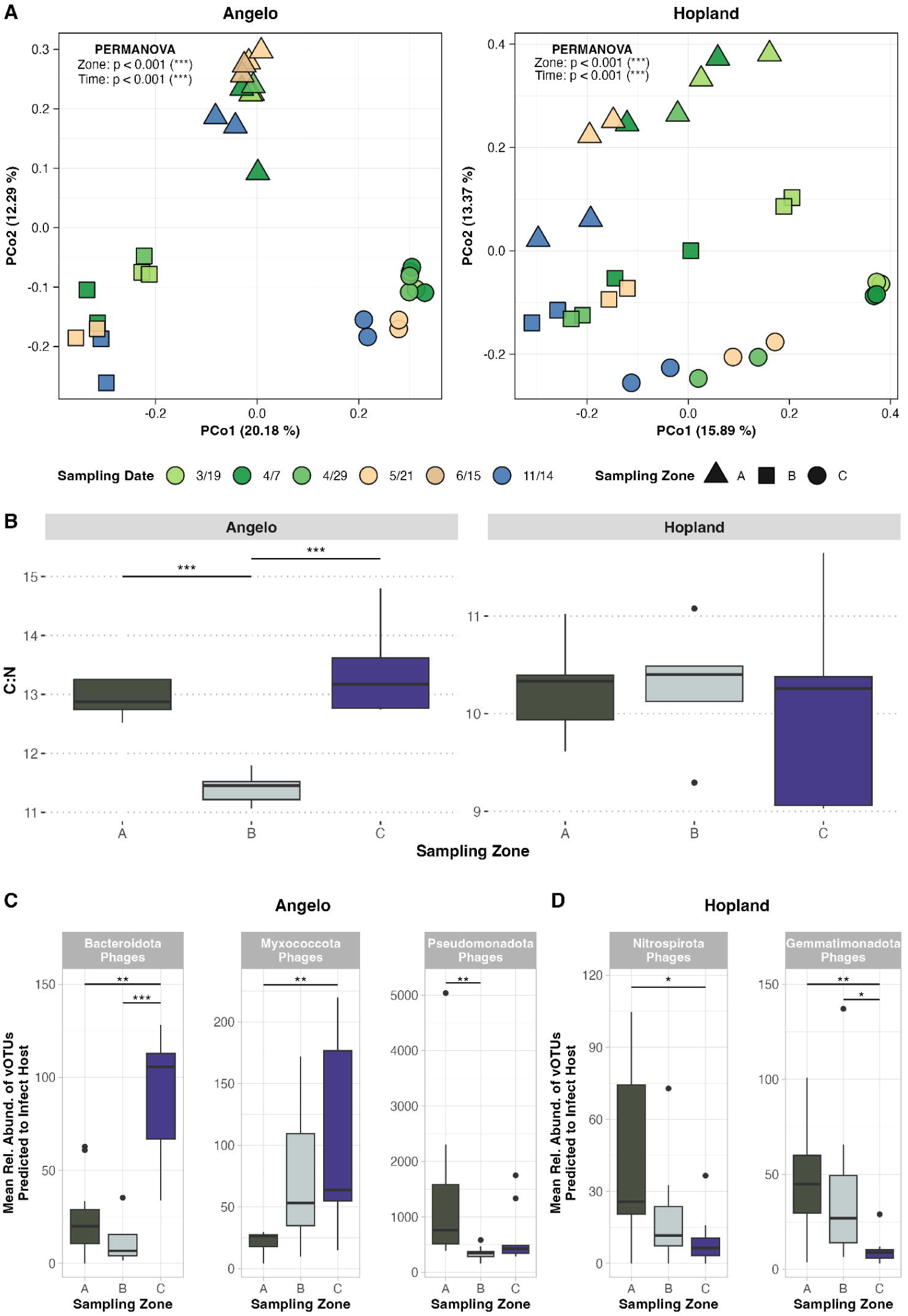
Horizontal spatial structuring controls viral communities recovered from viromes, soil edaphic properties, and abundances of viruses with predicted bacterial host phylum. **A)** Principal components analysis (PCoA) performed on Bray-Curtis dissimilarities of viral communities (vOTUs), with each point relating to one sample, faceted by site. Colors correspond to sampling time point and shapes denote sampling zone. Axis labels indicate the percentage of total variance explained. PERMANOVA statistics are reported in the top left of each plot. **B)** C:N ratio differences between sampling zones faceted by site. Significance between groups is marked by connecting lines with significance stars (*, p <0.05; **p <0.01; ***p<0.001), based on analysis of variance (ANOVA) and emmeans post-hoc test. Box boundaries correspond to 25th and 75th percentiles, and whiskers extend to ±1.5x the interquartile range. Boxplots are colored by sampling zone. **C, D)** Differences in mean relative abundances of viruses predicted to infect specific host phylum between sampling zones, summed per host phylum, faceted by host phylum in Angelo **(C)** and Hopland **(D)**. Box and whisker parameters, statistics, and colors are the same as in C.

Spatial differences at the >1 meter-scale were stronger at Angelo (Angelo R^2^ = 0.32, Hopland R^2^ = 0.24, **Table S1**), despite similar distances between samples at each site (Hopland mean = 15.5 m, max = 23.2 m; Angelo mean = 14.4 m, max = 19.9 m; **Figure 1B**), which may partly reflect environmental differences and co-variation with hosts. At Angelo, some edaphic parameters – total carbon, total nitrogen, and C:N ratio – differed significantly in zone B compared to zones A and C (**Figure 2B, Table S2**), while no environmental differences across zones were detected at Hopland (**Figure 2B, Figures S4A–B, Table S2**). Similarly, viruses grouped by predicted host phylum revealed nine host-associated viral groups whose abundances differed significantly among sampling zones at Angelo, including three of the five most abundant host groups in the viromic dataset (**Figure 2C, Figure S5A, Table S3-5**). In contrast, only two predicted host groupings showed zone-specific differences at Hopland (**Figure 2D, Figure S5B, Table S3-4**), and these viral host groupings were less abundant overall (**Table S5**), suggesting that they played relatively minor roles in the observed spatial structuring. Together, these findings point toward environmental and host filtering contributing to the stronger spatial structuring observed at Angelo than Hopland.

However, neither environmental nor host factors fully accounted for the spatial differences observed in both sites. At Angelo, no measured environmental variable or predicted host group was consistently different across all three sampling zones, and at Hopland, spatial structuring still occurred in the absence of significant environmental or predicted host gradients. These patterns are consistent with prior studies reporting strong spatial structuring within habitats [5, 25], including when host communities were directly measured [22, 23, 27], rather than predicted as in our study. Such observed spatial patterns are often interpreted as evidence for viral dispersal limitation. In our study, this explanation is plausible, but difficult to confirm because only four edaphic properties were measured, and host communities were not directly quantified, thus biotic and abiotic filtering could still be at play. Similarly, other unmeasured factors, such as historical contingency (e.g., legacy effects of past disturbances), microscale habitat heterogeneity (e.g., soil aggregates, microtopography, root distributions), biotic interactions (viral predation or competition), or stochastic processes could also contribute to the observed spatial structuring. To more clearly resolve the mechanisms underlying this spatial structuring, future field studies should broaden the suite of environmental measurements, directly characterize host communities, and incorporate fine-scale spatial sampling that captures microsite variability. Further, approaches that explicitly test dispersal, such as controlled soil microcosms or microfluidics, would be particularly valuable for determining whether movement constraints drive viral turnover across space.

### Temporal changes in surface viral communities were also significant, though secondary to spatial differences

Viral communities also differed significantly over time at both field sites, with turnover largely progressive over time, rather than showing periods of similarity between different times of year. While temporal structuring was apparent along the first PCoA axis at Hopland, it was masked by spatial variation at Angelo and only became evident along the third PCoA axis (**Figure 3A**). This likely reflects the fact that sampling zone accounted for more variation at Angelo compared to Hopland (30% vs. 24%), overshadowing temporal trends (**Table S1**). To better isolate the effects of time from sampling location, we examined temporal distance-decay relationships within each sampling zone. In this analysis, temporal decay was still stronger at Hopland (slope = 6 × 10^-4^) than at Angelo (slope = 4.9 × 10^-4^) (**Table S6**), and viral communities at the most distant timepoint comparisons were more dissimilar at Hopland (92% of Bray-Curtis sample comparison values >0.8) than at Angelo (all <0.8), suggesting greater temporal viral turnover for Hopland overall (**Figure 3B**).

**Figure 3:**
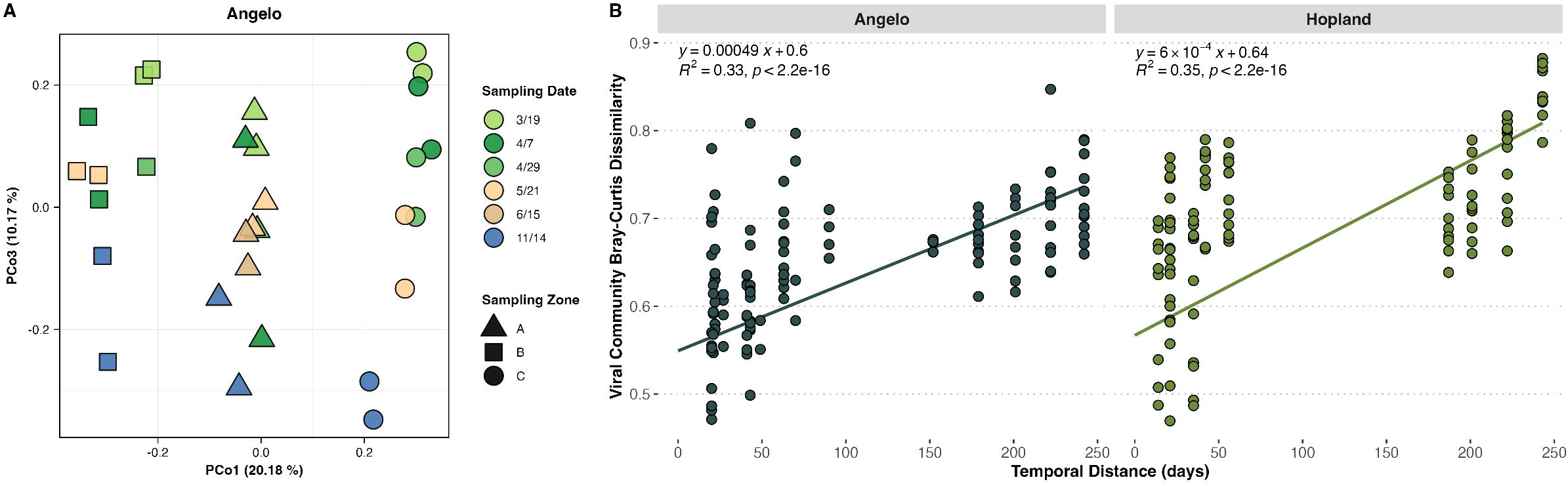
Progressive temporal structuring of viral communities derived from viromes. **A)** Principal components analysis (PCoA) performed on Bray-Curtis dissimilarities of viral communities (vOTUs) in Angelo, here displaying the first and third axis, which is where temporal structuring is observed. Each point relates to one sample, with color corresponding with sampling time point and shapes with sampling zone. Axis labels indicate the percentage of total variance explained. PERMANOVA statistics are reported in each plot. **B)** Temporal distance-decay curves, representing the relationship between community dissimilarity (Bray-Curtis) and change over time (days) in viral communities, faceted by site. Each point is a pairwise comparison between two samples, however, to assure spatial variation did not obscure temporal trends, samples comparisons were only made between samples taken from the same sampling zone (all three within zone comparisons plotted here). Trend lines display linear regressions, with associated formula and statistics reported in each facet.

The observed temporal trends could have been driven by changes in the soil environment over time, especially since sampling largely captured the three most distinct parts of the water year (**Figure 1D-E**). Linear regressions of total soil environment differences showed that the soil environment shifted more over time at Hopland (slope = 7.4 × 10^-3^) than at Angelo (slope = 5 × 10^-3^, **Figure S6A, Table S6**), and loess fits revealed that the soil environment at Angelo remained more similar across the most distant timepoints (**Figure S6B**), indicating weaker seasonal change. Underlying these patterns, gravimetric water content, pH, and total N all exhibited significant temporal distance-decay at Hopland, whereas only pH showed this trend at Angelo (**Figure S6C-D, Table S6**). While time has often generally proven to be a weaker explanatory force, relative to environmental condition or space, in both soil viral and microbial ecology [5, 13, 22, 30, 75, 76], one study in which viral temporal differences were more apparent than differences over space or depth included large seasonal shifts, specifically, transitions from snow-covered soils to peak plant growth [28]. Our results fit within this framework: temporal structuring in viral communities was more apparent at Hopland where environmental conditions changed more strongly through time. Taken together, the spatial and temporal results indicate that soil viral communities are organized by a layered hierarchy of controls, with spatial heterogeneity setting the dominant template and temporal change adding additional, but more moderate, structure. Extending temporal studies over longer periods would be valuable to determine whether environmental fluctuations shape viral communities (for example, through reoccurrences of seasonally driven viruses) or whether spatial and temporal interactions remain the primary drivers of community structure.

### Temporal variation outweighed spatial patterns in viral richness and abundance in surface soils

While viral community composition turned over in a progressive manner with time, viral richness exhibited non-linear and variable differences across our sampling timepoints. Overall, when considering time point as a categorical variable (as opposed to pairwise distance between days), viral richness differed significantly with time at both sites (p < 0.001, **Table S7**). Although richness was also significantly different between sampling zones at Angelo (p < 0.001, **Figure S7A, Table S7**), sampling zone explained less variation in Angelo viral richness (**Table S7**). Changes in richness over time followed different patterns at each site. At Angelo, richness remained relatively constant over the first three months, until the final timepoint (November, late wet-up), when it dipped to its lowest values (**Figure 4A**). A similar decline was observed during late wet-up at Hopland, but richness only dropped to levels comparable to timepoints during plant productivity. At Hopland, richness peaked during early dry-down (May), with values far exceeding those at Angelo during the same period, although most other timepoints were similar between sites (**Figure 4A**). Overall, the contrasting temporal trajectories at the two sites highlight that viral richness is not only sensitive to seasonal changes in soil moisture, with late wet-up emerging as a key period of decline, but also specific to local environmental conditions.

**Figure 4:**
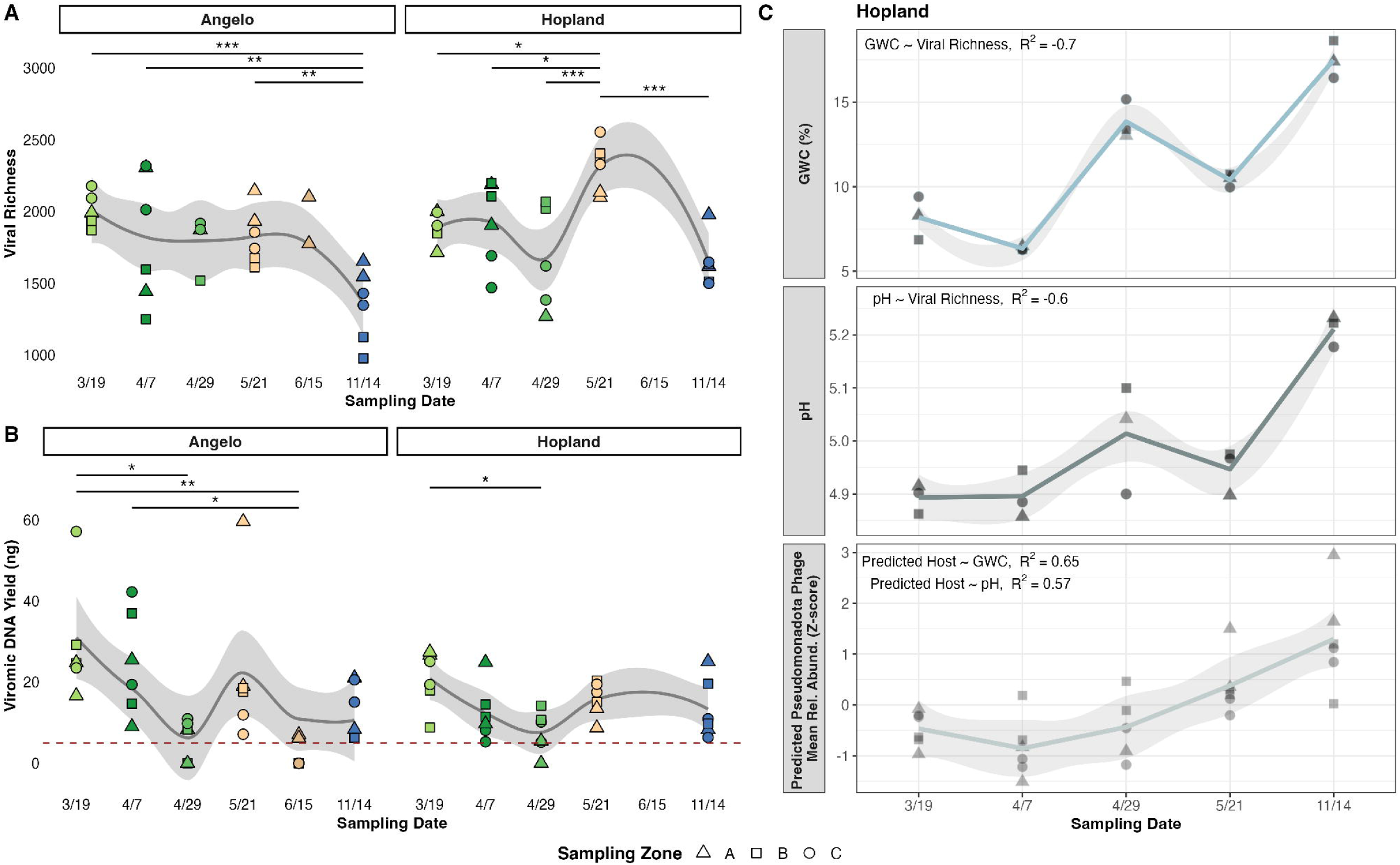
Non-linear temporal structuring of viral richness and viral DNA yields, underlain partially by soil edaphic and host properties in Hopland. **A)** Changes in viral community richness generated from total numbers of vOTUs over time, faceted by plot. Each point is one sample, with colors denoting sampling date and shapes denoting sampling zone. Trends lines report loess models. Significance between groups is marked by connecting lines with significance stars (*, p <0.05; **p <0.01, ***p<0.001), based on analysis of variance (ANOVA) and emmeans post-hoc test. **B)** Viromic DNA yields (a relative proxy for viral particle abundances) for each extracted virome over time, faceted site, with substitution of 0 ng for all viromes with yields that were below detection limits (detection limit of 0.5 ng, denoted by red dashed line). Statistics, colors and shapes are the same as in panel A. **C)** Changes in gravimetric water content (GWC, top panel - blue), pH (middle panel - green), and relative abundances of viruses predicted to infect *Pseudomonadota* (z-score transformed, bottom panel - gray) throughout time. Trend lines reflect fitted points predicted from loess models, with standard error (light gray) reported from these models. Spearman correlations between Hopland viral richness and GWC/pH, and viral *Pseudomonadota* abundance with GWC/pH, are reported at the top left of each plot. Shapes correspond with sampling zone.

Relative to richness trends, viromic DNA yields (a proxy for viral particle abundance) displayed more similar temporal patterns between sites (**Figure 4B**) and were only structured by time and not sampling zone (Angelo: p = 0.002, Hopland: p = 0.02, **Table S7**). At both sites, viromic DNA yields were highest during early plant productivity (March) and declined to a low point in late plant-productivity (April). Four of the six samples collected in late dry down (June) were below detection limits at Angelo, as were one (Hopland) and two (Angelo) samples collected in late plant productivity (April), suggesting low virion abundances [23]. Low moisture is thought to reduce the abundance of virions [3, 46], a pattern that matches the low-moisture conditions observed at Angelo during late dry down (**Figure S4C**). However, this relationship did not hold during late plant productivity, when soil moisture was higher than at other timepoints in the dataset (**Figure S4C-D**), yet DNA yields still declined. Our results largely mirror observations from a natural prairie over similar timescales, where viral abundances (measured by epifluorescence microscopy) peaked in March and May, with a dip in April [29], suggesting that factors beyond moisture (e.g. plant phenology) may influence virion abundances. Viral richness and DNA yields were positively correlated in each site, but these correlations were weak, and only slightly stronger in Hopland (Angelo R^2^ = 0.18, p = 0.02, Hopland R^2^ = 0.25, p = 0.0061, **Figure S7B, Table S6**).

These temporally fluctuating patterns in richness and viral DNA yields suggest an influence of the environment, such as moisture content. Thus, we compared richness and yield temporal trends with environmental variables and predicted hosts, retaining only moderate-to-strong Spearman correlations (|R^2^| ≥ 0.5). We used this approach because sampling times and depths differed for some measurements, so comparisons were based on similarity of temporal patterns rather than direct regressions. Under these criteria, only viral richness at Hopland showed qualifying correlations, as DNA yields at Hopland and both richness and yields at Angelo fell below the threshold. Hopland viral richness was negatively correlated with trends in both gravimetric water content (GWC, R^2^ = −0.7) and pH (R^2^ = −0.6, **Figure 4C, Table S8**) throughout time, indicating that richness tended to increase when soils were drier and more acidic. This trend is consistent with findings from one previously reported set of Hopland soil microcosms, where viral richness decreased after water addition [7], and with field studies reporting higher viral diversity in historically drier grassland ecosystems [47]. However, other Mediterranean grassland microcosm experiments, including some using Hopland soils, found increased viral diversity sustained for up to 10 days after rewetting [46]. It is possible that the decrease in viral richness with increased soil moisture observed in our study, which reflects *in situ* soil moisture dynamics rather than microcosm precipitation events, captured a broader suite of interacting biotic and abiotic processes that co-vary with moisture in the field. For example, grassland bacteria have been shown to respond and grow at different rates in response to soil moisture [42], and thus the reduction in overall viral richness could be attributed to host-specific replication of a subset of viruses, leading to temporary dominance of specific viral species. Alternatively, higher moisture could increase viral decay rates, as moisture can desorb viruses from soil surfaces [77], making them more available for degradation by other members of the soil food web through increased mixing [36, 78]. These mechanisms, individually or in combination, may contribute to the observed decrease in viral richness during wetter periods. Precipitation (and thus higher soil moisture) is also known to decrease pH in soils [79], which potentially resulted in the concurrent negative relationship between viral richness and pH. While a prior study found higher soil viral richness at more basic pH levels (e.g., pH 7.5 vs. 4.5) [19], our study focused on a much narrower and acidic range (pH 4.9–5.2) and scaled multiple timepoints that underwent fluctuations in moisture, indicating that temporally variable moisture-pH interactions, rather than differences in absolute pH levels across sites, may underlie the observed correlations with viral richness here.

Host responses are likely a key driver that links environmental variables to changes in viral dynamics, so we next sought to compare environmental variables to viral relative abundances grouped by predicted hosts. Hopland viruses predicted to infect the host phylum, *Pseudomonadota* were positively and strongly correlated with GWC and soil pH (**Figure 4C, Table S8**). This could indicate that specific *Pseudomonadota* host responses to the environment impacted a subset of viruses, whereas other hosts may have been less shaped by environmental drivers, preventing similar patterns from emerging in the broader viral community. *Pseudomonadota* are known to increase in abundance with increasing soil moisture [80, 81], which may help to explain the higher relative abundance of their predicted viruses in wetter Hopland soils. Together, these results suggest that viral richness and abundance vary temporally more than spatially, potentially influenced by environmental and host drivers, as was observed in the Hopland soils.

### Depth dominated viral community structure as measured in total metagenomes

To understand viral dynamics with depth and throughout additional parts of the water-year, we leveraged metagenomes collected from late dry-down (late April through July, compared to March through May for viromes) and early wet-up (September, relative to November for viromes) from three depths (0–10 cm, 20–30 cm, and 50–80 cm) (**Figure 1A**, [39]). Because total metagenomes are known to assemble substantially less soil viral diversity than viromes [22], we used the vOTUs assembled from the viromes as references for metagenomic read mapping to measure viral community composition in the metagenomes. At both sites, viral communities recovered from metagenomes were structured primarily by depth (PERMANOVA p < 0.001, Angelo R^2^ = 0.39, Hopland R^2^ = 0.29) and, to a lesser extent, sampling zone (PERMANOVA p < 0.001, Angelo R^2^ = 0.13, Hopland R^2^ = 0.10, **Figure 5A, Table S1**). At Hopland, time was also a marginally significant explanatory factor for viral community composition in metagenomes (R^2^ = 0.03, p = 0.041, **Table S1**). These communities were also weakly associated with total nitrogen at Angelo (R² = 0.03, p = 0.010) and pH at Hopland (R² = 0.03, p = 0.033). Because these environmental variables covaried with depth (**Figure S8A-B, Table S10**), we regressed them against depth and used residuals in generalized additive models (GAMs) to isolate their independent effects. This analysis revealed that over 700 vOTUs (∼10% of total, **Table S9**) remained significantly associated with total nitrogen even after accounting for depth, suggesting nitrogen plays a role independent of vertical distribution of DNA viruses. In contrast, only two vOTUs were independently associated with pH, indicating a tighter coupling between pH and depth.

**Figure 5:**
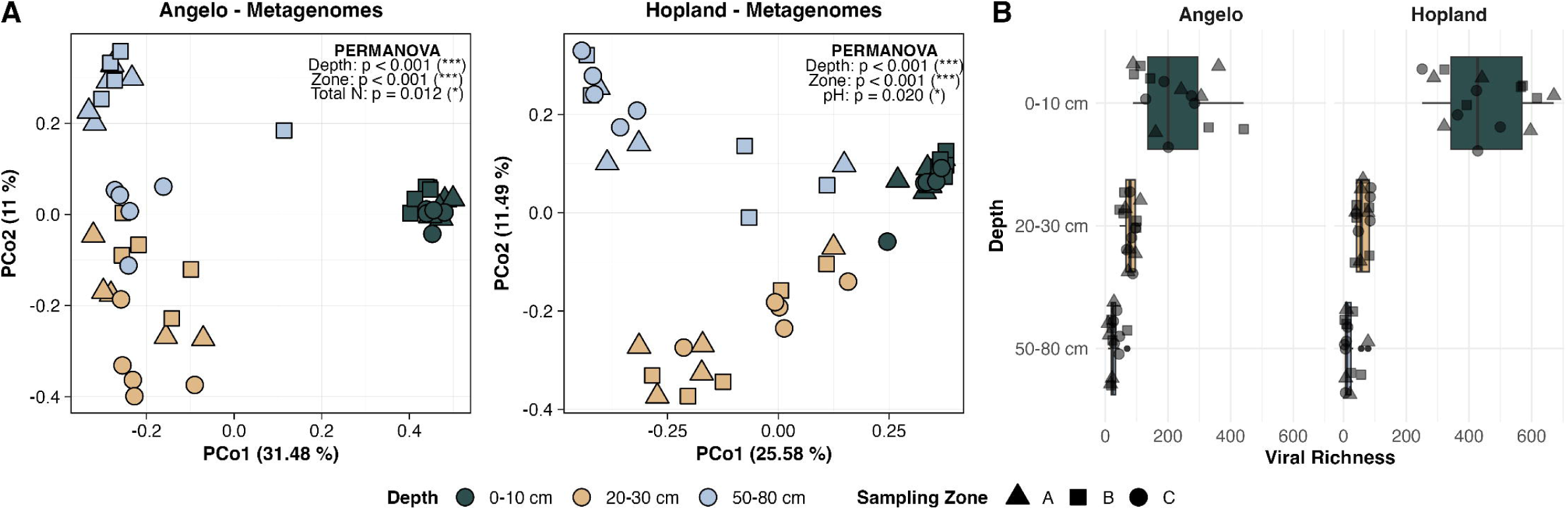
The effects of depth and sampling zone on viral communities derived from metagenomic read-mapping. **A)** Principal Components Analysis (PCoA) performed on Bray-Curtis dissimilarities of viral communities (vOTUs) derived from metagenomic read-mapping, faceted by site. Each point is one sample, where colors correspond with depth and shapes with sampling zone. Axis labels indicate the percentage of total variance explained. Statistics from PERMANOVA analyses are reported in each plot. **B)** Changes in viral community richness across soil depths, generated from total counts of vOTUs, faceted by site. Each point is one sample, where sampling zone is designated by shape. Boxplot colors correspond with soil depth, with box boundaries and whiskers the same as panel B.

Both soil environment and host distributions may have contributed to the strong spatial and depth-based structuring we observed for viral communities recovered from metagenomes. At both sites, pH increased and total carbon and nitrogen decreased with depth, with pH also varying by sampling zone (**Figure S8A–B, Table S10**). Prokaryotic communities from these same samples showed similar spatial patterns to the viral communities, and were structured by depth and sampling zone with little influence of time [39]. The prokaryotic communities were also shaped by pH [39], echoing the patterns observed in the Hopland viral communities. Together these results suggest that both environmental and host gradients through space could have influenced viral communities, leading to the strong horizontal and vertical spatial structuring observed. Interestingly, unlike previous studies where viruses often show stronger spatial structuring than bacteria [13, 22, 23], here both communities displayed comparable spatial patterns, suggesting shared ecological controls across soil depths and sites.

### Viral richness decreased with depth and aligned with host distributions

Viral community richness declined significantly with depth at both Angelo and Hopland sites (**Figure 5C**), with intermediate (20–30 cm) and deep (50–80 cm) soils exhibiting significantly lower richness compared to surface soils (0–10 cm; linear mixed-effects model, **Table S11**), with no effects of horizontal spatial variation (**Table S11**). While the observed vertical patterns could be partially due to methodological biases – since viromes were constructed from surface soils, diversity measured in metagenomes via read mapping to vOTUs assembled from viromes could be inflated at these shallow depths – similar depth-related declines in bacterial diversity were observed from these samples [39]. Decreased microbial diversity with depth is a common phenomenon [82–84], and is typically attributed to environmental constraints such as reduced nutrients, lower oxygen, and cooler temperatures [85–87]. These factors not only limit bacterial diversity but may also restrict viral richness by reducing host availability and activity. Supporting this, Penev et al. [39] reported that some bacterial phyla, particularly *Actinomycetota*, *Pseudomonadota*, and *Acidobacteriota*, were most active (e.g., relative isotope incorporation) in deeper soils in the metagenomes leveraged in our study. We observed that the diversity of vOTUs predicted to infect these host taxa was also significantly higher in deep soils (*Actinomycetota* and *Acidobacteriota* at Angelo, *Actinomycetota* and *Pseudomonadota* at Hopland, **Figure S9A&B, Table S12**), reinforcing the interpretation that host abundance was an important filter shaping viral diversity.

## Conclusions

Our findings reveal that spatial structuring consistently outweighed temporal variation in shaping soil viral communities. When depth was assessed, it emerged as the strongest structuring factor, surpassing both horizontal spatial and temporal influences. These patterns were partially attributable to edaphic differences in the samples and predicted and directly measured host distributions, suggesting a potential role for environmental and biotic filtering in shaping viral community composition. However, these factors did not fully explain the observed variation, indicating that unmeasured environmental variables, dispersal limitation, or other biotic interactions likely contributed substantially to viral community spatial structuring. In contrast, viral richness and DNA yields were closely tied to seasonal changes, suggesting that the total number of viral species and overall virion abundance respond more strongly to environmental pulses, like moisture. Together, these results highlight the need for explicit testing of soil viral spatial dynamics at finer scales linked to a broader range of environmental and host-associated measurements (e.g., host activity, metabolite gradients) that may drive spatial heterogeneity. Additionally, longer-term, inter-annual temporal studies are needed to capture viral dynamics that may not manifest over a single seasonal cycle.

## Acknowledgements

We would like to thank the Hopland Research and Extension Center and Angelo Coast Range Reserve staff for facilitating site access and providing sampling support. We are also grateful to members of the DOE Science Focus Area (SFA) Microbes Persist team, LLNL staff scientists, and members of the Banfield, Firestone and Pett-Ridge labs who helped with soil sampling and processing, and to Christina Wistrom and the Oxford Tract Greenhouse team at UC Berkeley for use of the facility for soil processing. We thank all members of the Emerson Lab, specifically Christian Santos-Medellin, Anneliek ter Horst, Sara Geonczy, Grant Gogul and Luke Hillary, for valuable discussions on viral ecology and data analysis support.

## Funding Sources

Funding for this work was provided by the U.S. Department of Energy, Office of Biological and Environmental Research (DOE-BER), Genomic Science Program, LLNL ‘Microbes Persist’ Soil Microbiome Scientific Focus Area (grant SCW1632 (to JP-R) and a UC Davis subaward (to JBE)). Work at Lawrence Livermore National Laboratory was conducted under the auspices of the U.S. DOE under contract DE-AC52-07NA27344.

## Data Availability

All raw sequences have been deposited in the NCBI Sequence Read Archive under the BioProject accession PRJNA1264504. The database of dereplicated vOTUs is available at 10.5281/zenodo.15512610. All scripts for data processing are available at https://github.com/janefudyma/WaterYearViromes.

## Notes

### Competing Interest Statement

The authors have declared no competing interest.

