## Supplemental Figures for "Spatial and depth structuring predominate over temporal variation in Mediterranean grassland soil viral communities"

### Supplementary Figures for manuscript “Spatial and depth structuring predominate over temporal variation in Mediterranean grassland soil viral communities”

Jane D. Fudyma, Petar Penev, Katerina Estera-Molina, Jordan Hoff, Steven J. Blazewicz, Jennifer Pett-Ridge, Joanne B. Emerson

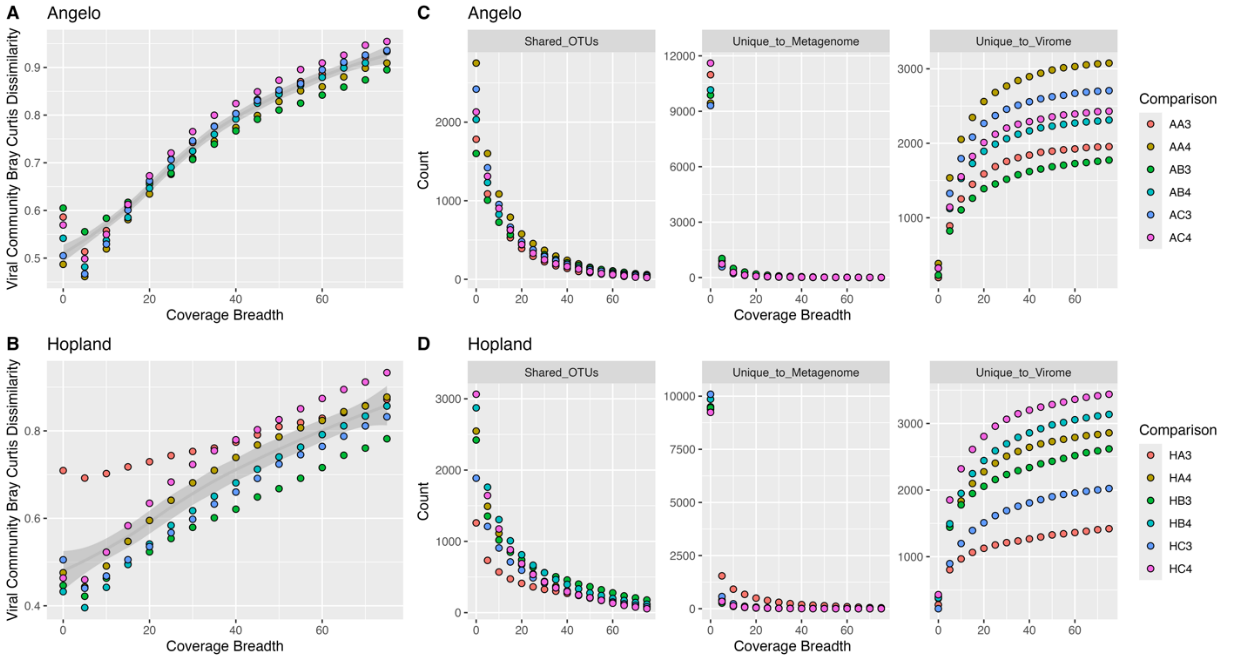


**Figure S1.** To identify a suitable coverage breadth threshold for quantifying vOTU abundances from metagenomic data, metagenomic reads were mapped to a vOTU database using a range of coverage breadth cutoffs (from 75%, the standard for vOTU recovery from viromes, down to 0%, in 5% increments). For each threshold, two comparisons were performed: (A-B) Bray-Curtis dissimilarity between viral communities detected in metagenomes and paired viromes from the same samples, assessing compositional similarity across thresholds; and (C-D) presence-absence overlap of vOTUs between metagenomes and viromes, categorized as shared, metagenome-exclusive, or virome-exclusive. These comparisons were conducted separately for each study site to support selection of a threshold that balances stringency and vOTU recovery for downstream analyses. Legend: the second letter denotes the sampling zone, and the number denotes the sampling time, where samples were only compared for two timepoints.

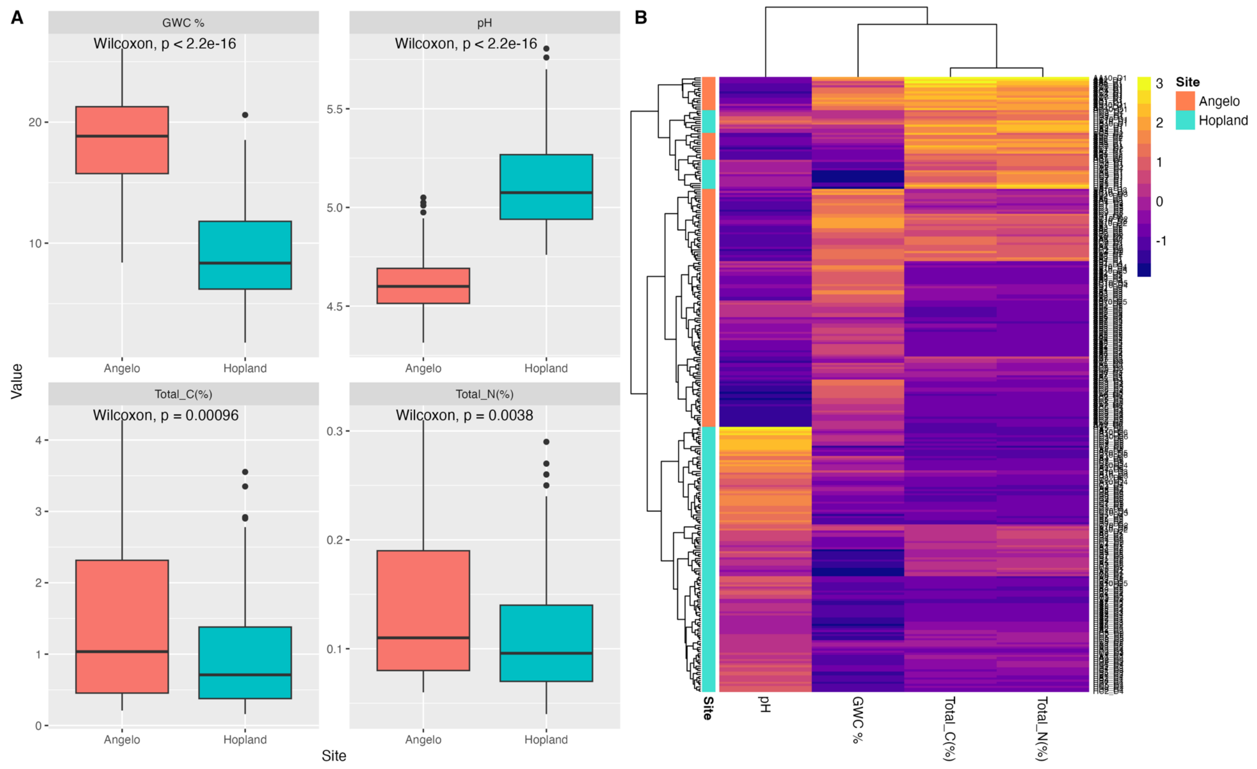


**Figure S2:** A) Soil environmental measurements by site. Box boundaries correspond to 25th and 75th percentiles, and whiskers extend to ±1.5x the interquartile range. Colors denote site sampled. P-values from Wilcoxon tests are displayed at the top of each plot. B) Heat map of z-score transformed soil edaphic variables by both site, with colors in tree corresponding to each site.

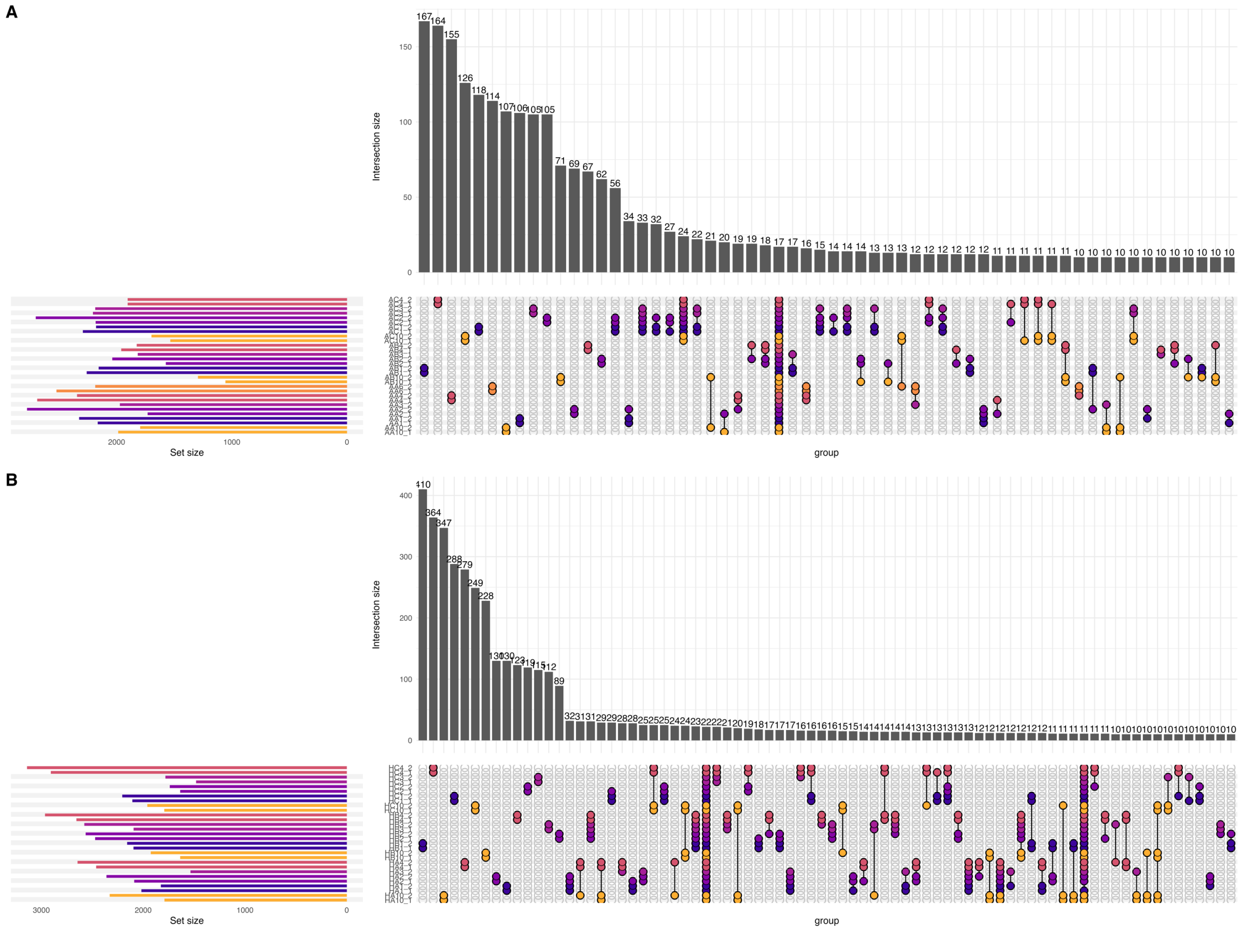


**Figure S3.** UpSet plot of shared vOTUs in each sample for Angelo (A) and Hopland (B), based on presence-absence data derived from read mapping to vOTUs. Colored dots indicate the sample(s) in which a given set of vOTUs was detected, and connecting lines between dots indicate vOTUs shared between samples. Sample IDs follow the format SZT_#, where: the first letter indicates the site (A = Angelo, H = Hopland), the second letter denotes the sampling zone, the third character (number) indicates the sampling timepoint, and the final digit identifies the sub-replicate (1 or 2).

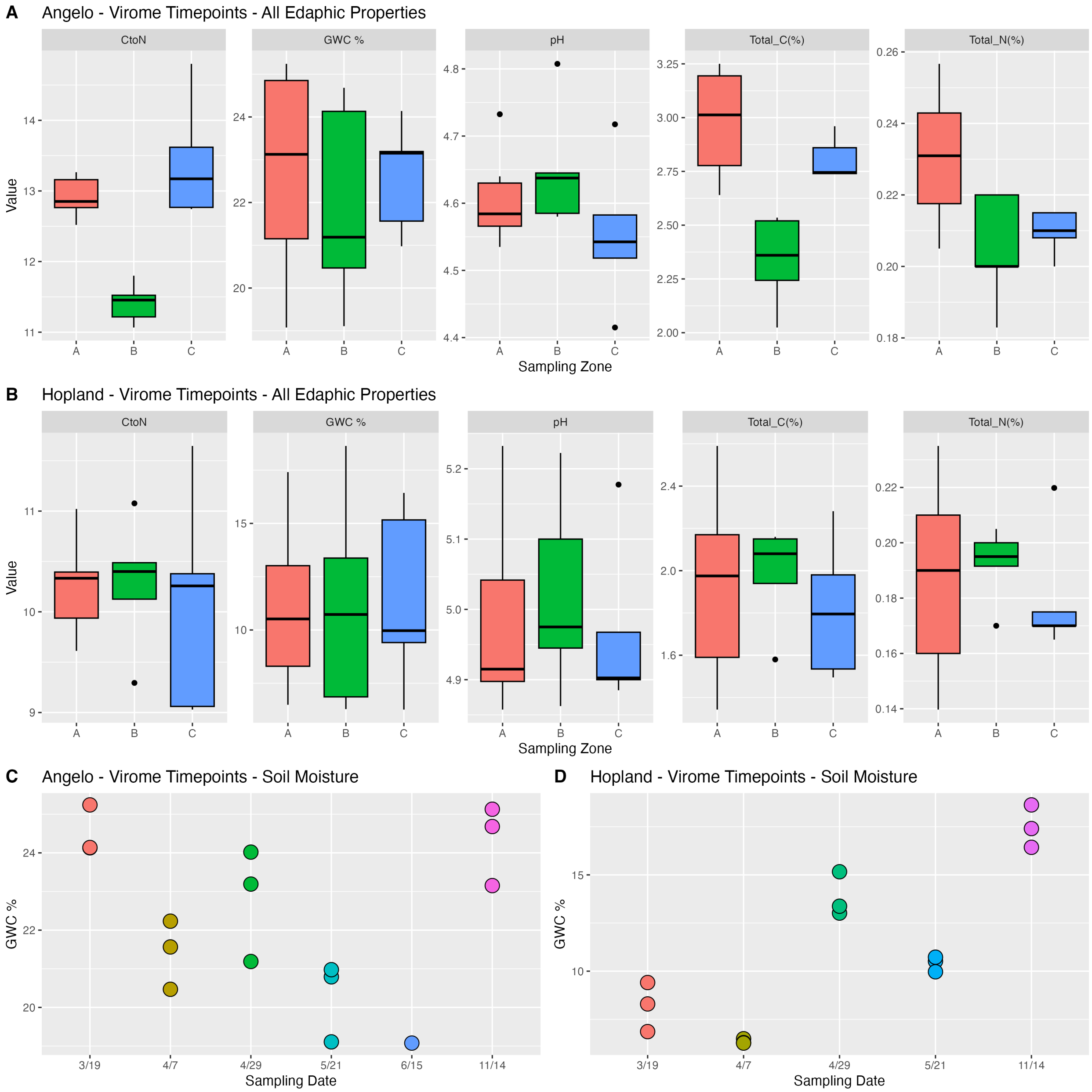


**Fig S4:** Differences in measured soil edaphic properties by sampling zone in **A)** Angelo and **B)** Hopland for the timepoints where viromes were processed. Box boundaries correspond to 25th and 75th percentiles, and whiskers extend to ±1.5x the interquartile range. Colors denote each sampling zone sampled. ANOVA or Chi-Squared with emmeans post-hoc tests for significant comparisons are reported with bars and significance stars (*, $p < 0.05$; **, $p < 0.01$, ***, $p < 0.001$). Gravimetric Water Content (GWC %) changes throughout time in **C)** Angelo and **D)** Hopland for the timepoints that viromes were processed. Colors correspond with sampling date.

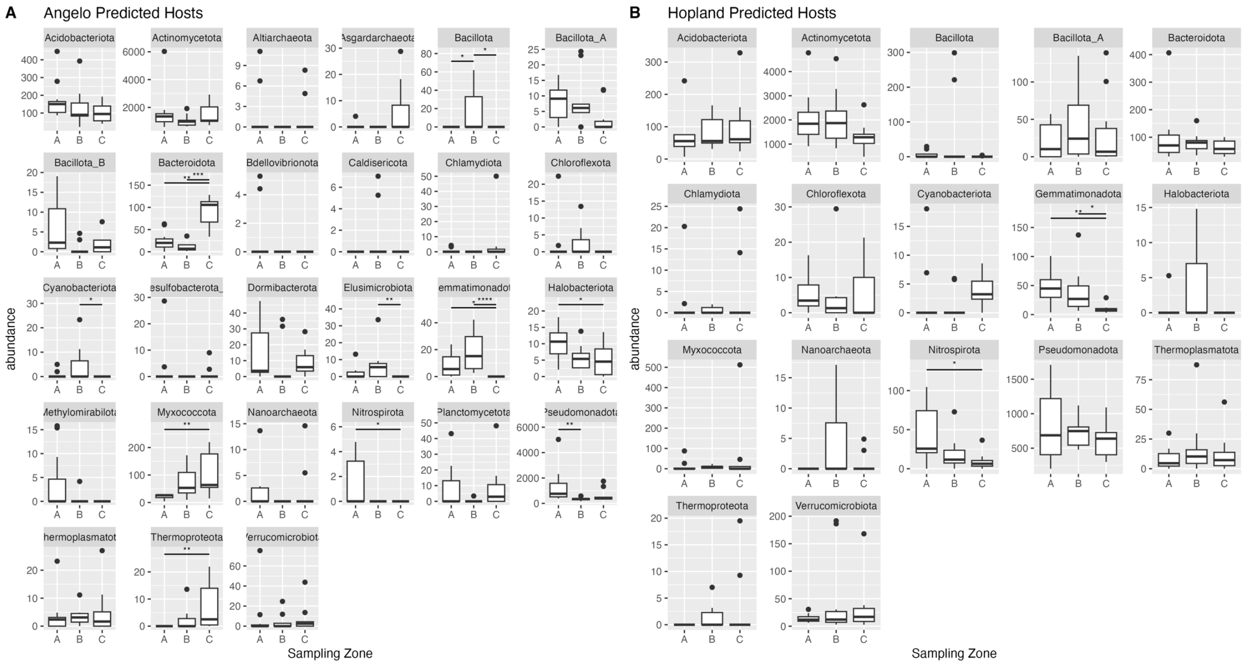


**Fig S5**: Differences between sampling zones of viral abundances predicted to infect different hosts in each site (panel A – Angleo, panel B – Hopland). Box boundaries correspond to 25th and 75th percentiles, and whiskers extend to ±1.5x the interquartile range. ANOVA with emmeans post-hoc tests for significant comparisons are reported with bars and significance stars (*, p <0.05; **, p <0.01, ***, p < 0.001).

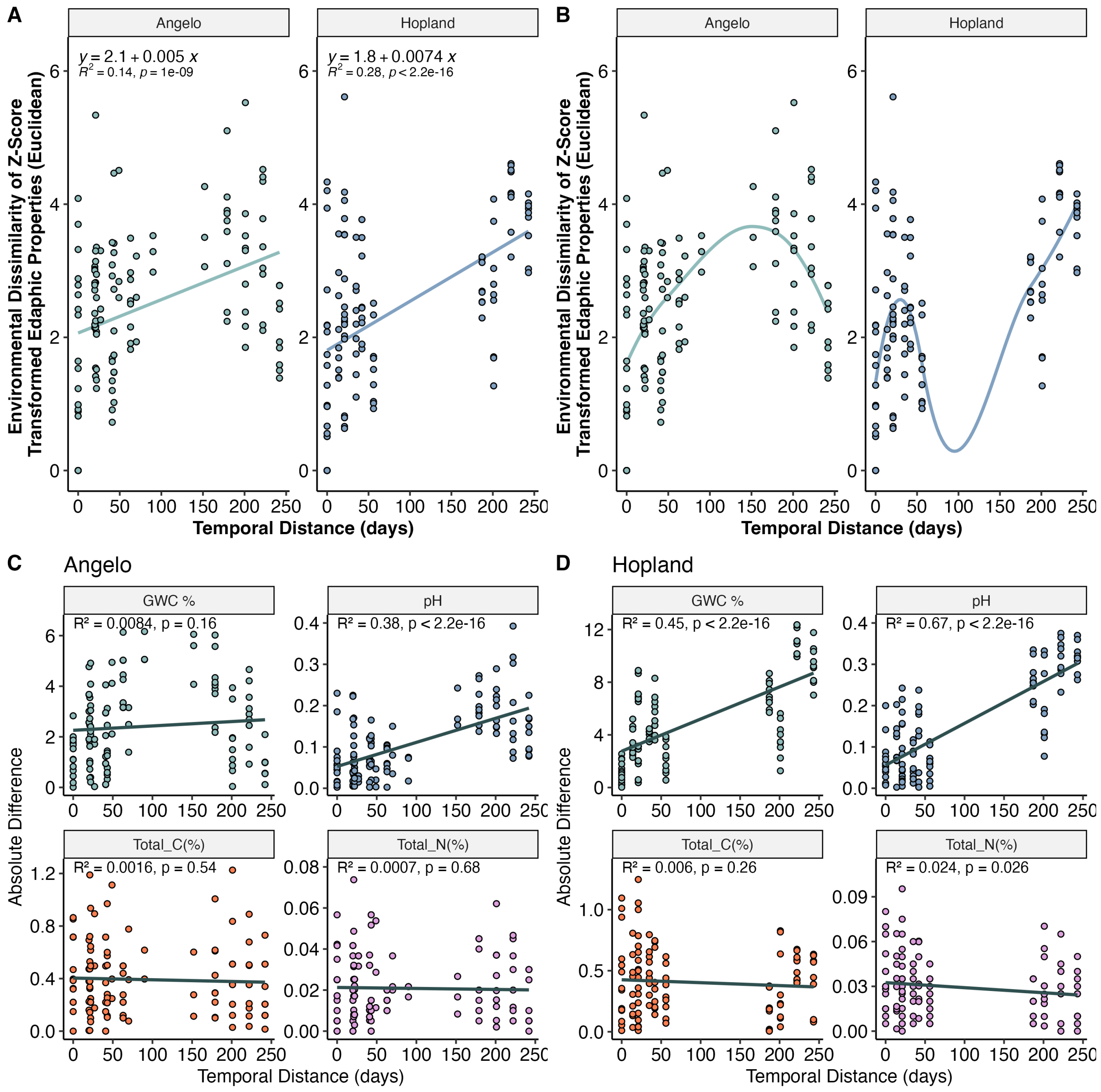


**Fig S6.** Temporal distance-decay curves, representing the relationship between the change over time (days) and either the total environment (z-score transformed – panels A&B) or specific environmental parameters (panels C&D). Each point is a pairwise comparison between two samples. Trend lines display linear regressions (panels A, C &D) or loess models (panel B), with associated statistics reported in each facet.

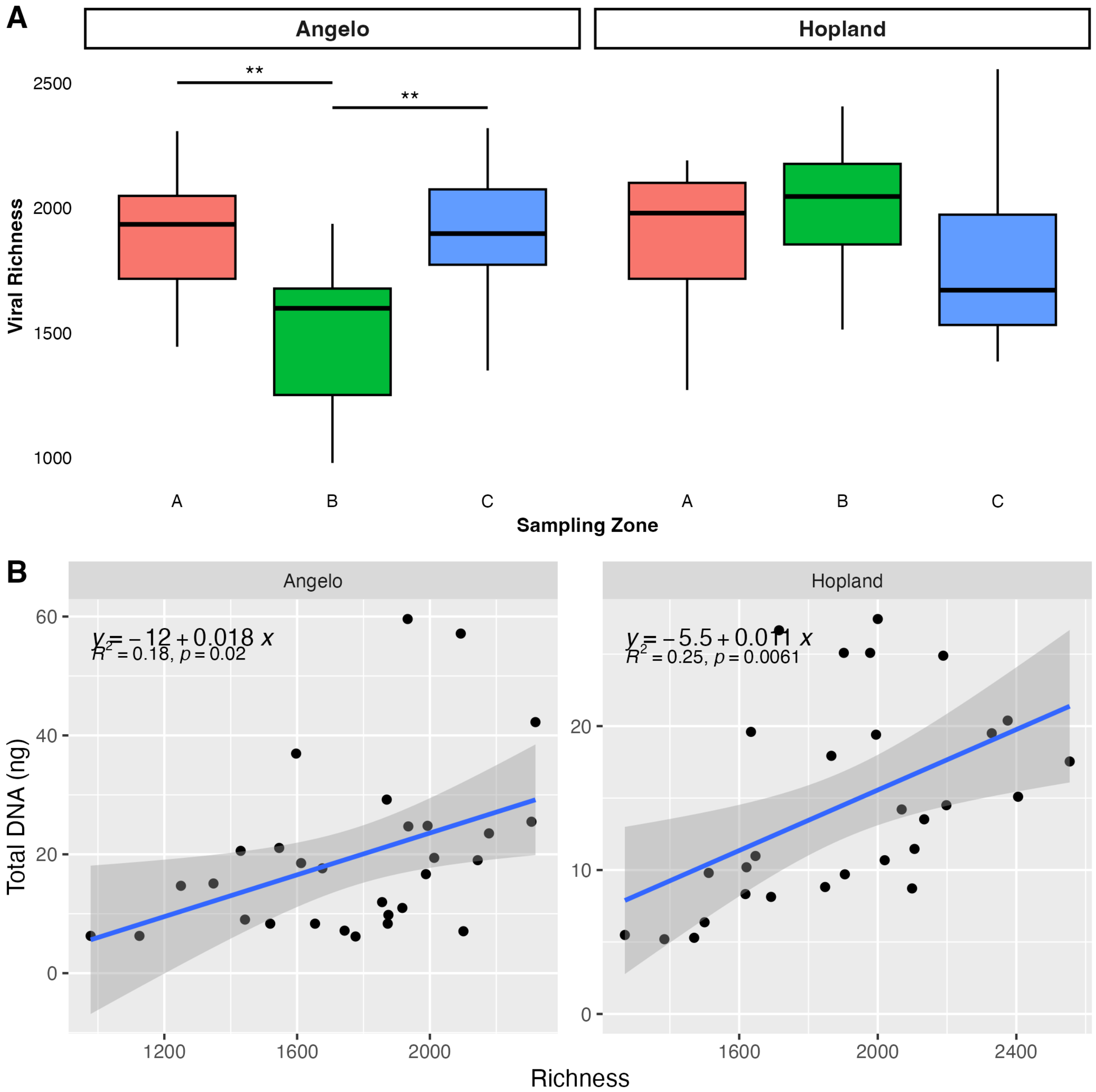


**Fig S7: A)** Differences in viral richness by sampling zone, faceted by site. Box boundaries correspond to 25th and 75th percentiles, and whiskers extend to ±1.5x the interquartile range. Colors correspond with sampling zone. ANOVA with emmeans post-hoc tests for significant comparisons are reported with bars and significance stars (*, $p < 0.05$; **, $p < 0.01$, ***, $p < 0.001$). **B)** Correlations between viral richness and total viral DNA, faceted by site. Linear regressions are fit to each plot, with corresponding equation, $R^2$, and p-values reported.

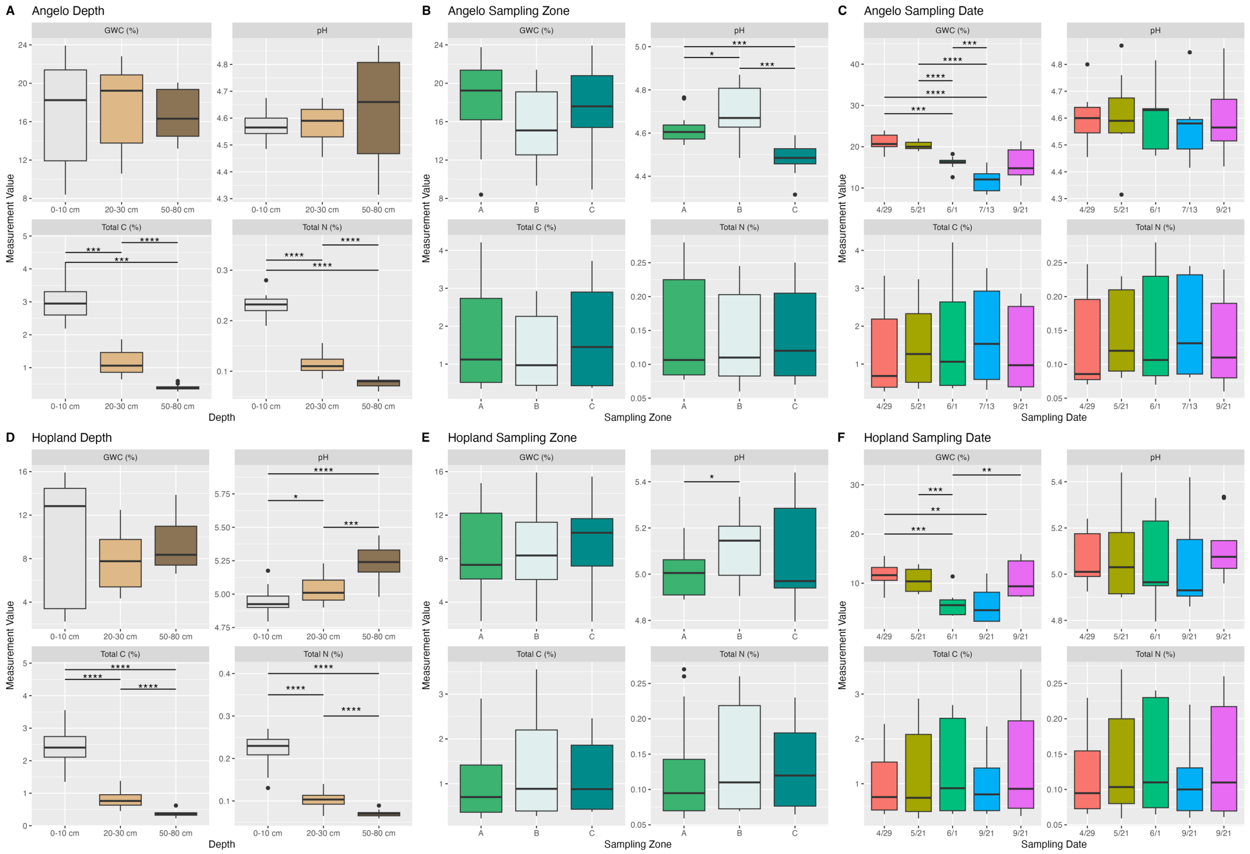


**Fig S8:** Differences in measured soil edaphic properties by depth (A,D), sampling zone (Rep, B, E) and sampling date (C-F). Top panels correspond with Angelo, bottom panels correspond with Hopland. Box boundaries correspond to 25th and 75th percentiles, and whiskers extend to ±1.5x the interquartile range. Colors denote different depths, zones, or timepoints. ANOVA with emmeans post-hoc tests for significant comparisons are reported with bars and significance stars (*, p <0.05; **, p <0.01, ***, p < 0.001).

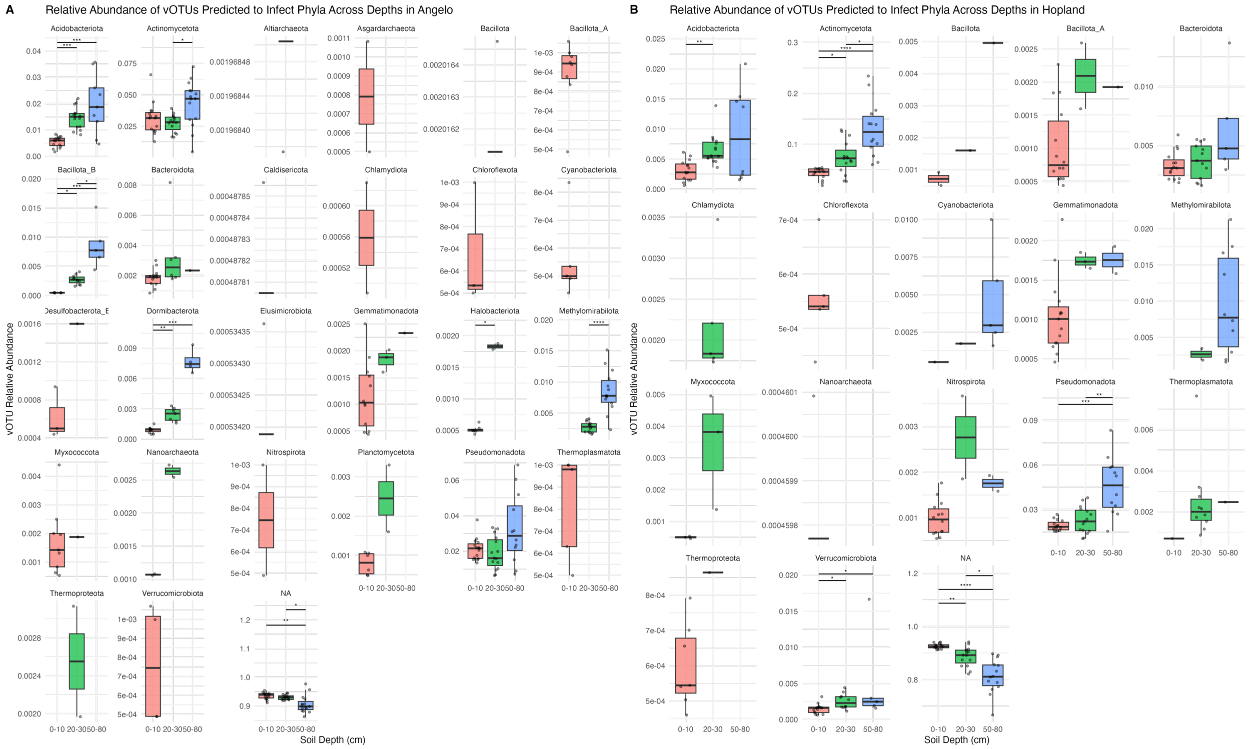


**Fig S9**: Differences between depths of viral relative abundances predicted to infect host phylum in each site, generated from metagenomic read mapping to virome vOTUs in A) Angelo and B) Hopland. Box boundaries correspond to 25th and 75th percentiles, and whiskers extend to ±1.5x the interquartile range. Colors represent different depths. ANOVA with emmeans post-hoc tests for significant comparisons are reported with bars and significance stars (*, $p < 0.05$; **, $p < 0.01$, ***, $p < 0.001$).
